# Anthropogenic impacts on threatened species erode functional diversity in turtles and crocodilians

**DOI:** 10.1101/2022.03.11.483822

**Authors:** R.C. Rodríguez-Caro, E. Graciá, S.P. Blomberg, H. Cayuela, M. Grace, C.P. Carmona, H.A. Pérez-Mendoza, A. Giménez, R. Salguero-Gómez

## Abstract

The Anthropocene is tightly associated with a drastic loss of species worldwide and the disappearance of their key ecosystem functions. The on-going reduction in ecosystem functionality is driven by global and local threats. The orders Testudines (turtles and tortoises) and Crocodilia (crocodiles, alligators, and gharials) contain numerous threatened, long-lived species for which their functional diversity and potential erosion by anthropogenic impacts remains unknown. Here, we examine 259 (69%) of the existing 375 species of Testudines and Crocodilia, quantifying their life history strategies (*i.e*., schedules of survival, development, and reproduction) from open-access data on their demography, ancestry, and threats. We find that the loss of functional diversity in simulated extinction scenarios of threatened species is greater than expected by chance. Moreover, the effects of unsustainable local consumption, diseases, and pollution are associated with specific functional strategies. In contrast, climate change, habitat disturbance, and global trade affect all species independent of their life history strategy. Importantly, the loss of functional diversity for threatened species by habitat disturbance is twice that for all other threats. Our findings highlight the importance of conservation programmes focused on preserving the functional diversity of life history strategies jointly with the phylogenetic representativity of these highly threatened groups.

## INTRODUCTION

Human impacts, such as habitat loss, climate change, pollution, poaching, and unsustainable trade, impose major threats to the persistence of species worldwide (Maxwell et al., 2016). Indeed, modern human activities have drastically accelerated species extinctions by three orders of magnitude (Pimm et al., 1995). As such, effective conservation measures are urgently needed to bend the curve of biodiversity loss (Díaz et al., 2019).

The on-going biodiversity loss is resulting in the erosion of functional diversity, key to ecological function (Oliver et al., 2015; Carmona et al., 2021; Toussaint et al., 2021). A robust way to characterise diversity in functional traits across a specific clade is via quantifying life history strategies, *i*.*e*. trade-offs between an individual’s investment in survival, development, and reproduction driven by limited resources and physiological constraints (Stearns, 1999; Salguero-Gómez et al., 2016a; Cernansky, 2017; Healy et al., 2019; Capdevila et al., 2020). In this context, different species can possess similar life history trait values, resulting in functional redundancy (Naeem, 1998; *e*.*g*., the Texas tortoises, *Gopherus berlandieri*, and the common snapping turtle, *Chelydra serpentina*, achieve similar longevities; Germano, 1994), or display dissimilar functionality, giving rise to functional dispersion (Elmqvist et al., 2003; *e*.*g*., the loggerhead sea turtle, *Caretta caretta*, reproduces later than other Testudines of similar size, such as the Aldabra giant tortoise, *Aldabrachelys gigantea*; Scott et al., 2012). Understanding how redundant or dispersed life history traits are and how sensitive these are to anthropogenic threats worldwide is critical for conservation (Chichorro et al. 2019).

Evaluating the risk of extinction of long-lived species is important for an accurate understanding and preservation of their functional diversity. Critically, the extinction of long-lived species can be delayed due to their long generation times (Gaillard et al., 1989; Jackson and Sax, 2010; Salguero-Gómez et al., 2016a; Capdevila et al. 2020). Thus, environmental effects typically manifest at much longer time scales in long-lived species than in short-lived ones (Gibbs and Amato, 2000; Kuussaari et al., 2009). Perhaps not surprisingly, two of the most threatened and long-lived groups are the order Testudines (tortoises, turtles, and sea turtles) and Crocodilia (crocodiles, alligators, and gharials) (Rhodin et al., 2018; Colston et al., 2020; Cox et al., 2022). In fact, Testudines and Crocodilia contain some of the highest proportions of threatened species across vertebrates (57.9% and 50%, respectively; Cox et al., 2022; UICN, 2020). Moreover, the conservation status of Testudines and Crocodilia could be even worse, since 25% of them lack reliable data to accurately identify their IUCN status (Rhodin et al., 2018).

Here, we characterise the life history strategies of 69% of Testudines and Crocodilia species –259 out of 375 extant species– to identify how the impacts of different human-led global (climate change, global wildlife trade, and emergent diseases) and local threats (habitat disturbances, unsustainable local consumption, and pollution) may alter their functional diversity. Using demographic and phylogenetic open-access data, we quantify life history strategies to estimate their functional diversity. We then simulate different extinction scenarios according to their extinction risk and the main threats that affect each species. We hypothesise that: (H1) the loss of functional diversity due to the extinction of threatened species will be lower than expected by chance. This is so because the life history strategies across testudines and crocodilians are expected to show high functional redundancy, since most of them are long-lived species with delayed maturity (Healy et al., 2019); (H2) human threats (*e*.*g*., global trade, unsustainable local consumption, disease) will differentially erode the functional diversity of these species (Ripple et al., 2017). For instance, some evidence exists that tortoise and turtle populations may be declining due to habitat disturbance, unsustainable local consumption, and international pet trade collection (Stanford et al., 2020). In contrast, threats to the functional diversity of these species due to global climate change may *a priori* not seem more intense than local threats (Cox et al., 2022). However, a fundamental knowledge about the consequences of climate change in long-lived species, such as Testudines and Crocodilia, is still to emerge. Filling in this knowledge gap is the primary motivation of this research.

## RESULTS

### High functional redundancy

Two dominant axes of life history traits describe most of the functional diversity of Testudines and Crocodilia. Using life history trait data from 236 species of turtles and tortoises (67% of the extant species), and 23 species of crocodilians (85%) from multiple sources, we characterise their life history strategies via a phylogenetic principal component analysis (*p*PCA; Table S1) accounting for adult body mass (Table S2). To address gaps in the dataset (See Appendix 1), we performed a multiple phylogenetic-trait imputation for missing values (see Materials and Methods). The two dominant axes of life history strategies encompass 62% of the total functional variation (Fig. 1). These axes correspond to (i) the ‘fast–slow continuum’ (Sæther, 1987; Stearns, 1999; Oli, 2004; Bielby et al 2007), which separates species with short maximum lifespans, like some freshwater turtles (*e*.*g*., 4.3 years for the Alabama red-bellied turtle, *Pseudemys alabamensis*), to species with long maximum lifespans such as terrestrial tortoises (*e*.*g*. 177 years for the Pinzon Island giant tortoise, *Chelonoidis ephippium*); and (ii) the ‘reproductive strategy’ (Gaillard et al., 1989; Salguero-Gómez et al., 2016a; Hughes, 2017; Paniw et al. 2018; Capdevila et al. 2020), defined by a trade-off between clutch frequency *vs*. reproductive output, with species with high numbers of clutches per year but lower clutch size per clutch at one end (*e*.*g*., 1.2 eggs per clutch and three clutches per year, as in the Madagascan flat-tailed tortoise, *Pyxis planicauda*), and extremely fecund species at the other end, having high clutch sizes but few clutches per year (*e*.*g*., 91.5 eggs per clutch and one clutch per year, as in the South American River Turtle, *Podocnemis expansa*).

**Fig. 1.**
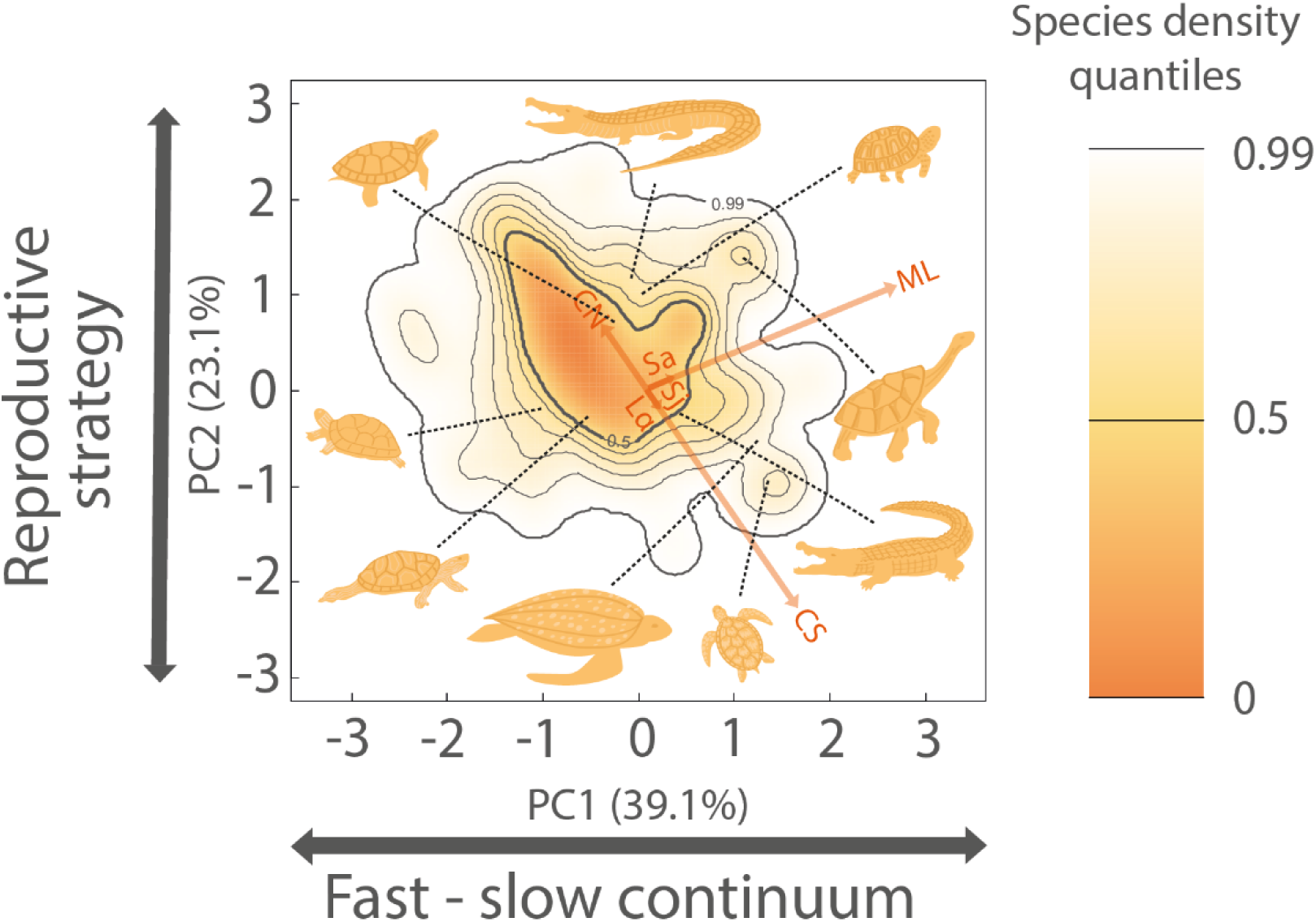
The global functional spectra of the life history strategies of Testudines and Crocodilia is described by two dominant axes of life history variation: (PC1) fast-slow continuum and (PC2) reproductive strategies. The probabilistic species distribution in this space is defined by the two first principal components axes (PC1 = 39.1% and PC2 = 23.1% of variance explained) of a phylogenetically-corrected principal component analysis (*p*PCA), where adult body mass has also been corrected. The phylogenetic signal is Pagel’s *λ* = 0.674 ± 0.030 (sd). The life history traits are: adult survival (*Sa*), juvenile survival (*Sj*), maximum lifespan (*ML*), age at sexual maturity (*Lα*), mean of number of clutches per year (*CN*), clutch size (*CS*). Arrows indicate the direction and weighting of each trait in the *p*PCA. The colour gradient (orange, yellow, and white) depicts the density of species in the defined space, where orange corresponds to more densely populated areas). Thick contour lines indicate the 0.5 and 0.99 quantiles, and thinner ones indicate quantiles 0.6, 0.7, 0.8, and 0.9. The species silhouettes correspond to (starting at the top left and moving clockwise): painted turtle (*Chrysemys picta*), American alligator (*Alligator mississippiensis*), desert tortoise (*Gopherus agassizii*), Hood Island giant tortoise (*Chelonoidis hoodensis*), Orinoco crocodile (*Crocodylus intermedius*), loggerhead sea turtle (*Caretta caretta*), leatherback sea turtle (*Dermochelys coriacea*), northern map turtle (*Graptemys geographica*), and Western Caspian turtle (*Mauremys rivulata*).

Examining the functional spectra of the life history traits of testudines and crocodilians reveals a high degree of functional redundancy. We use a trait probability density (TPD) approach considering the two-dimensional functional space (Fig. 1) to identify peaks (areas of high density) and valleys (low density) of species diversity (Carmona et al., 2016; Toussaint et al., 2021). The examined reptile species tend to congregate around a single hotspot with median values of maximum lifespan and high number of clutches per year with lower clutch sizes. This hotspot of life history strategies includes 50% of the 259 species and represents 13.2% of the total spectrum (Fig. 1), in agreement with a recent analyses examining 6,567 species of reptiles (12.3% of the spectrum; Carmona et al., 2021). Freshwater turtles, such as the eastern long-necked turtle, *Chelodina longicollis*, (max. lifespan 37 years, 13.9 eggs/clutch, two clutches/yr) and crocodiles, such as the mugger crocodile *Crocodylus palustris* (max. lifespan 31.5 years, 28.75 eggs/clutch, two clutches/yr), are located in that hotspot of functional diversity. Sea turtles are represented in a valley of functional diversity (bottom right of Fig. 1), which is characterised by species with higher clutch sizes, few clutches per year, and higher maximum lifespan (*e*.*g. Caretta caretta:* max. lifespan 76 years, 115 eggs/clutch, 1.29 clutches/yr). In contrast, terrestrial tortoises are represented at the top right of Fig. 1, with their typically shorter clutch sizes, high clutches per year, and longer maximum lifespans (*e*.*g*., the Hood Island giant tortoise *Chelonoidis hoodensis*: max. lifespan 177 years, 10 eggs/clutch, three clutches/yr).

### Unique strategies may become extinct

The extinction of threatened species of testudines and crocodilians would result in the loss of a quarter of the global functional diversity of these taxa. To estimate the effect of the potential extinctions of threatened species on the existing amount of functional diversity in these groups, we use the IUCN Red List categories of our species and simulate an accumulative loss of threatened species (*i*.*e*., in the first scenario “-CR”, we remove the *Critically Endangered* species only; in the “-EN” scenario, we remove Endangered and Critically Endangered species, and so on; see Materials and Methods). If all threatened species disappeared (*Critically Endangered, Endangered*, and *Vulnerable* species; “-VU” scenario, Fig. 2A), the functional diversity would decrease by 26.8%.

**Fig. 2.**
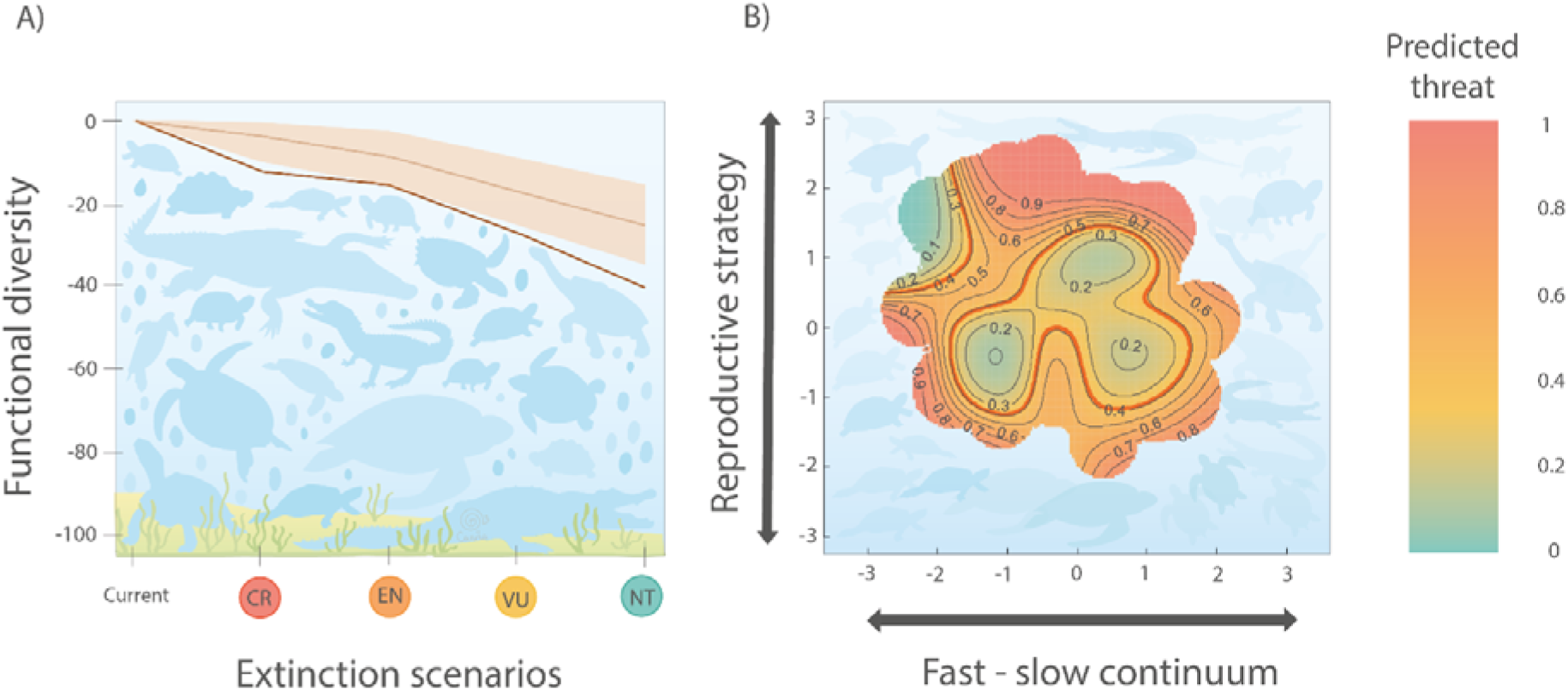
The life history strategies of testudines and crocodilians predict their vulnerability to extinction due to human threats. **A**) Loss of functional diversity under extinction scenarios of threatened species. CR: Critically Endangered, EN: Endangered, VU: Vulnerable; NT: Near Threatened. The loss of functional diversity is expressed as a percentage of the current functional diversity in the life history strategies of species of the order Testudines and Crocodilia (Fig. 1). We simulate the loss of functional diversity by removing all species from the IUCN category in a progressive manner, by degree of “endangerment”, removing first the species with a higher risk of extinction (*i*.*e*. -CR), then continuing progressively to remove species from the categories with lower threatened risks (-EN scenario: removing CR and EN species; - VU scenario: removing CR, EN & VU; -NT scenario: removing CR, EN, VU & NT). This loss of diversity is represented by a black line. For each scenario, we compare the loss of functional diversity with 999 iterations of a null model where threatened species are randomly selected among the 259 species in the analyses. The 999 randomisations are represented by a blue polygon defined by the 5^th^ and 95^th^ percentiles of functional diversity loss, with the blue line representing the 50^th^ percentile; **B**) Risk of extinction in the functional space of Testudines and Crocodilia species. Probability of species being classified as high-risk of extinction (*i*.*e*., CR & EN IUCN status) according to GAMs (with binomial distribution) using the position of each species in the two-dimensional functional space as predictors. Green tones indicate lower risk of extinction, whereas red tones indicate higher risk of extinction. *P* value of the GAM model is 0.03 and χ^*2*^ is 28.59. The red contour lines indicate the average threat probability (proportion of species classified as threatened in the group). Here, we consider only the 214 out of the 259 available species of testudines and crocodilians for which the threat status is known.

The loss of functional diversity due to the extinction of threatened species is greater than expected by chance. We compare the potential loss of functional diversity, considering the number of threatened species in each extinction scenario, with a simulated loss of functional diversity where the identity of the threatened species is randomised within the pool of species. The resulting loss of functional diversity due to the extinction of threatened species is greater than 95% of randomisations for most scenarios (Fig. 2A). In the “-CR” scenario, the loss of functional diversity (-12.67%) is four times greater than in randomisations (-3.82% [-0.4%, -9.69%] 5% and 95% CI). This finding suggests that most of the Critically Endangered species of testudines and crocodilians show unique life history strategies, such as the pancake tortoise (*Malacochersus tornieri* max. lifespan 25.9 years, one egg/clutch, one clutch/yr).

Unique or less redundant species strategies of testudines and crocodilians are more vulnerable to vanish following the extinction of threatened species. To identify differences in extinction risk between the life history strategies, we map extinction risk in the functional trait spectra via generalized additive models (GAMs), using the IUCN Red List category of each species (Carmona et al. 2021). The position of each species along the trait spectra is significantly correlated with its extinction risk when considering high-risk species as Critically Endangered and Endangered (*p* = 0.039; Fig. 2B). Using this grouping, high-risk species show distinct positions in the life history functional space, with species with lower functional redundancy located around the periphery and likely being more threatened. Only species located at the centre of the functional space and those with short maximum lifespan and smaller clutch sizes but with greater number of clutches per year show a lower probability of being threatened (Fig. 2B).

### Habitat disturbance as the main threat

The life history strategies of Testudines and Crocodilia can predict vulnerability to extinction due to three –local consumption, disease, and pollution– of the six analysed threats (Fig. 3). To explore the differential effects of global and local threats on the functional spectra of testudines and crocodilians, we analyse the probability of being affected by each threat separately using GAMs (more results in Fig. S1). Local consumption (Fig. 3A) is associated with increased extinction risk in species with higher clutch size such as sea turtles (*e*.*g*., *Caretta caretta*), and higher maximum lifespans, such as the radiated tortoise (*Astrochelys radiata*) or the Orinoco crocodile (*Crocodylus intermedius*). In contrast, diseases are associated to threaten species with slow life histories (Fig. 3B), such as the desert tortoise (*Gopherus agassizii*). Finally, pollution primarily is associated to species with higher reproductive output (*i*.*e*., higher clutch sizes), such as freshwater turtles like *Chelydra serpentina* or crocodilians like the saltwater crocodile, *Crocodylus porosus*, (Fig. 3C).

**Fig. 3.**
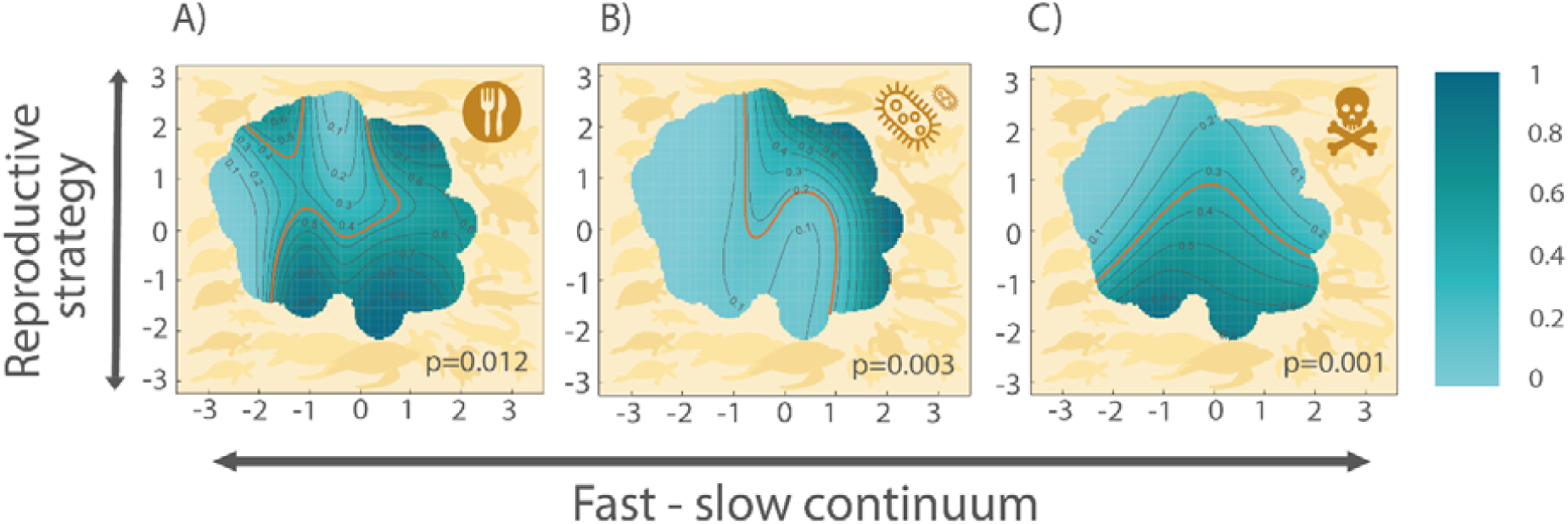
The life history strategies of Testudines and Crocodilia predict the vulnerability to extinction due to local consumption, disease, and pollution. Other risks, such as climate change, global trade, or habitat disturbance, do not show a significant relation with the examined species’ life history strategies (Fig. S1). Probability of species being affected by: **A**) local consumption, **B**) disease, and **C**) pollution, according to GAMs (with binomial distribution) using the position of species in the two-dimensional functional space (Fig. 1) as predictors. Light tones indicate lower risk of extinction due to each threat, whereas dark tones indicate higher risk. *P* values associated with each GAM are shown at the top-right corner of each panel. The red contour lines indicate the average threat probability. Here, we consider only species whose threats are known (n = 251 species).

Among all examined threats, the extinction of species affected by habitat disturbance results in the worst loss of functional diversity, especially in the northern hemisphere (Fig 4A). To estimate the effect of specific threats on the loss of functional diversity, we simulate the extinction of species affected by each threat to calculate the percentage of loss of functional diversity (Fig. 4B). We compare the potential loss of functional diversity by each threat, with a simulated scenario where the identity of the species affected by each threat is randomised within the pool of species. Most of the effects of loss of functional diversity due to the six threats considered are greater than expected by random extinctions, especially for climate change, whose effect is beyond the 95% percentile for random simulations (Fig. 4B). Only habitat disturbance and diseases show similar or lower values of loss of functional diversity than expected by random simulations (Fig 4B).

**Fig. 4.**
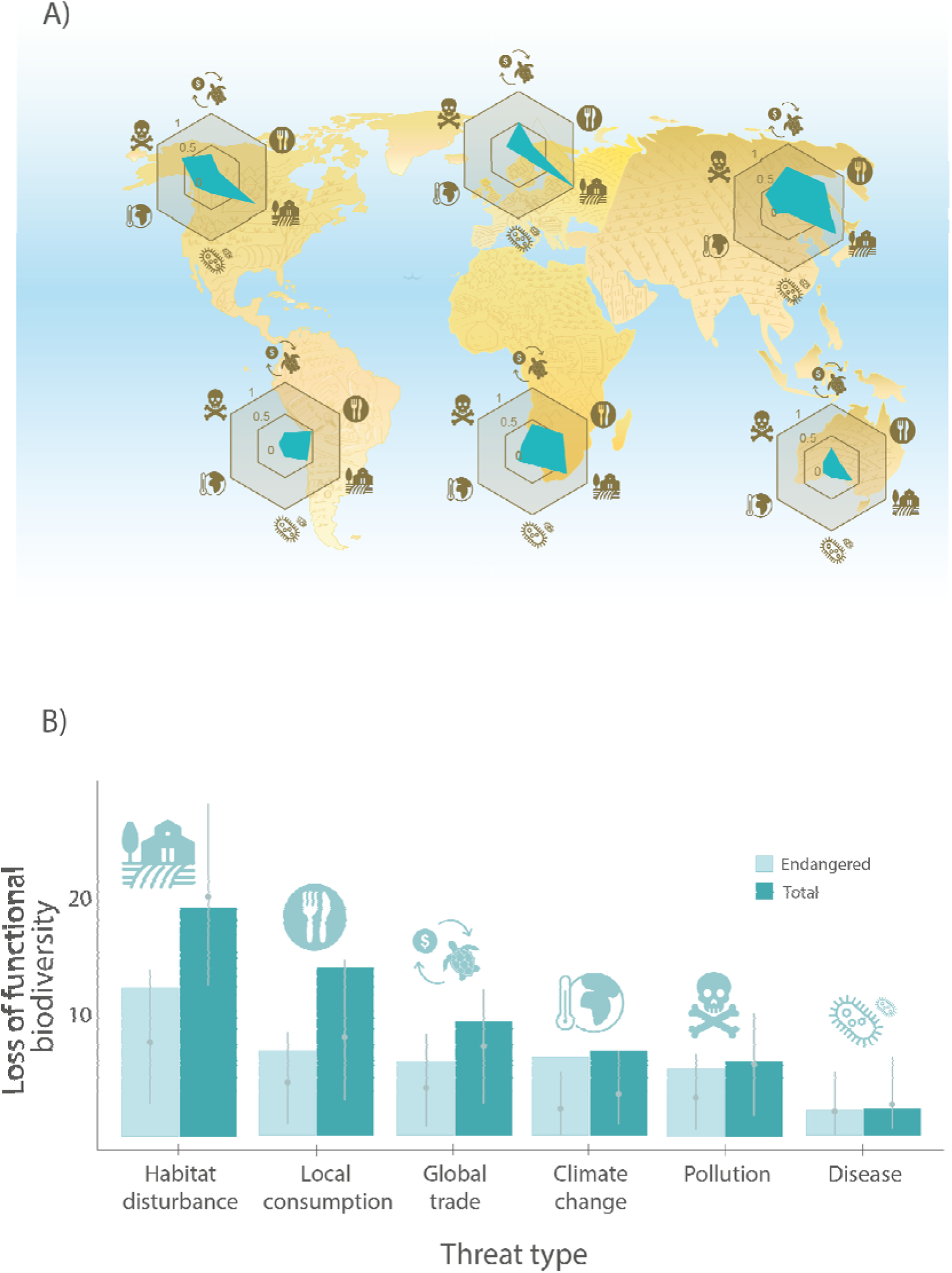
Habitat disturbance is the main threat for the functional diversity, especially in the north hemisphere. **A**) Ratio of species affected by each threat type per continent. The ratio is only estimated for species whose threats are known at the continent level (North America, n = 65; Europe, n = 4; Asia, n = 66; South America, n = 57; Africa, n = 39; Oceania, n = 14). **B**) Simulated loss of functional diversity of Testudines and Crocodilia under extinction scenarios by threat. The loss of functional diversity is expressed as a percentage of the total spectra of functional diversity of species, as detailed in Fig. 1. We simulate the loss of functional diversity by removing all species affected by specific threats, in light blue colour removing only threatened species (*i*.*e*. Critically Endangered [CR], Endangered [EN] & Vulnerable [VU] as per the IUCN Red List) and in dark blue removing all the species affected (threatened or not). For each scenario, we compare the loss of functional diversity with 999 iterations of a null model where the same number of species were randomly selected among all 251 species. The 999 randomisations are represented for each threat as a grey dot for the 50^th^ percentile, with grey whiskers representing the 5^th^ and 95^th^ percentiles.

## DISCUSSION

Here, we demonstrate that threatened species of turtles and crocodilians show great deal of functional dispersion. Toussaint et al. (2021), exploring 5,686 species of reptiles, found a loss of functional diversity linked to the extinction of the threatened species corresponding to less than 10%. Here, focussing on Testudines and Crocodilia, for which much information has been missing to date (Conde et al., 2019), we find twice the loss of functional diversity. By linking this information to their conservation status, we demonstrate that global and local threats are exposing the functional diversity of these already highly threatened taxonomic groups to great peril. Using the largest collection of trait data to date for turtles and crocodilians, we find that the threatened species show unique life history strategies. Losing these species due to extinction will result in a greater reduction of functional diversity and potentially severe ecological consequences (Violle et al., 2017) due to the ecosystem services they provide, such as the high values of biomass contribution of turtles (*e*.*g*., red-eared slider turtle, *Trachemys scripta*, Lovich et al., 2018), mineral cycling and bioaccumulation (*e*.*g*., *Chelydra serpentine*; Hebert et al., 1993), or seed dispersal (*e*.*g*., Galapagos tortoises, *Chelonoidis spp*; Blake et al., 2012).

Local disturbances such as habitat disturbance and unsustainable local consumption are the main threats for the functional diversity of the studied species. Habitat disturbances affect most species of turtles and crocodilians, especially in North America and Europe (Fig. 4A), independent of their life history strategies. Therefore, habitat protection for these species is key to preserve the functional diversity (Lourenço-de-Moraes et al., 2021). In fact, Mittermeier et al. (2015) highlighted that the protection of only 16 hotspot of turtle biodiversity would successfully encompass 83% of these species. On the other hand, we find that the second local threat for the functional diversity of Testudines and Crocodiles is unsustainable local consumption. Our results predict the extinction of species with higher rates of clutch size (*e*.*g*., such as sea turtles, whose eggs are consumed in high quantities; Madrigal-Ballestero et al., 2013) and long-lived species (linked with bigger size, which typically provides more meat; Donlan et al., 2010). Unsustainable local consumption of turtles is one of the most important threats in Asia, South America, and Africa (Fig. 4A). For example, in China, one million turtle farms are the primary purchasers of turtles caught in the wild, including several threatened species (Shi et al., 2007)

Global threats such as unsustainable or illegal international trade and climate change can also have important effects in reducing the functional diversity of Testudines and Crocodilia. The global trade of these species is the third most important threat for their functional diversity, affecting tortoises and turtles mainly for their use as pets (Luiselli et al., 2016) and crocodilians for their skin (Thorbjarnarson, 1999; Platt and Thorbjarnarson, 2000). On the other hand, although climate change affects fewer species than other threats, turtles and crocodilians with unique life history strategies could disappear according to our projections (Azanza-Ricardo et al., 2017). Indeed, the consequences of climate change for these species are found worldwide, including extreme droughts by compromising the fitness of the populations (Poiani and Johnson, 1991; Rodríguez-Caro et al., 2021), flood nesting sites by rising sea levels (Pike et al., 2015), coral bleaching due to increasing in temperatures, affecting feeding areas of sea turtles (Chaloupka et al., 2008) and the unbalance of sex-ratios in populations of temperature sex-dependent species (Janzen, 1994), such as the threatened leatherback turtle (*Dermochelys coriacea*; Tomillo et al., 2014).

Our study does not capture all human activities with impacts on species’ ecological strategies. As such, we deem our findings to be conservative. For instance, although we have summarised the breadth of testudines and crocodilians’ threats to extinction, threat interactions, which remain largely unknown in these taxonomic groups, have been reported to have synergistic effects on the loss of species, as was described for mammals (Romero-Muñoz et al., 2020) and birds (Chen et al., 2015). Moreover, examining the ecological functions of each species in higher resolution, by examining their diets or habitat breadth *(i*.*e*., generalist–specialist), could improve the available information of ecological function (Cooke et al., 2019).

Protecting threatened species with unique life history strategies is key to conserve their functional diversity (Violle et al., 2016) and ecosystem services they provide (Lovich et al., 2018). Our findings support previous studies that suggest that conservation policies should consider together the taxonomic and functional dimensions of biodiversity (Carmona et al., 2021) to implement targeted and more effective conservation actions in the context of a global biodiversity crisis (Toussaint et al., 2021). The IUCN Red List of Threatened Species may include information about population trends, distribution area, threats, and abundance. The inclusion of functional diversity could greatly aid international, national, and local efforts to preserve functionally unique species, and recently, new indexes have been developed to include some aspects of functional diversity. The IUCN Green Status of Species (Grace et al. 2021; IUCN 2021), a new part of the Red List assessment, quantifies species recovery and explicitly includes ecological function (Akçakaya et al., 2020). Incorporating the functional diversity of life history strategies in conservation assessments constitutes a promising approach to help prioritise conservation efforts to maintain high levels of functional diversity against current and future threats.

## MATERIALS AND METHODS

### Life history traits

To describe the life cycle of species in the orders Testudines and Crocodilia, we used life history trait data. Life history traits quantify how the life cycle of a species and its underlying vital rates —survival, development, reproduction— have been shaped by natural selection to optimise its performance (Roff, 1992). We obtained such traits from published literature, open access databases, and via direct researcher contributions (Cayuela et al. 2019). In some cases, life history trait information was derived from the demographic information of different databases and datasets. These data were obtained from the COMADRE Animal Matrix Database (Salguero□Gómez et al. 2016b), DATLife Database (2021), Amniote Life History Database (Myhrvold et al. 2015), and the reviews by Allen et al. (2017), Pfaller et al. (2018) and Reinke et al. (under review) (more information about data selection in Appendix S2).

To quantify species’ life history strategies, we selected six life history traits that encompass detailed information regarding the timing, intensity, frequency, and duration of demographic processes across the life cycle of any species (Stearns, 1992). The life history traits chosen were adult survival (*Sa*), juvenile survival (*Sj*), maximum lifespan (*ML*), age at sexual maturity (*Lα*), mean number of clutches per year (*CN*), and clutch size (*CS*; Table S1). These traits are well-established metrics in comparative demographic analyses, and their emerging syndromes typically explain high variance (∼75%) in life history strategies of multicellular organisms (Bielby et al., 2007; Carmona et al., 2021; Gaillard et al. 1989, 2005; Paniw et al., 2018; Salguero□Gómez et al. 2016a; Stearns 1992). The data obtained from the aforementioned sources encompass 259 species with at least one of these six traits (more information about data collection/estimation in Appendix S2).

### Phylogeny and data imputation

To identify and quantify the potential role of evolution in shaping our species’ life history strategies, we performed phylogenetic comparative analyses. We used a species□level phylogenetic tree recently published for Testudines and Crocodilia (Colston et al. 2020) to both impute the missing data and account for the effect of evolutionary constraints in the observed life history strategies. Briefly, to build the phylogenetic tree, 14 mitochondrial loci and six nuclear loci were sampled for 357 species of extant or recently extinct turtle and tortoises, and the 27 species described crocodilian species (Rhodin et al. 2017). Colston et al. (2020) computed Maximum Likelihood trees with RaxML (Stamatakis 2014) and used taxonomic imputation to fill the gaps of 17 out of 384 (<5%) species in the tree. Further details on the methods for tree construction are available in Colston et al. (2020). We selected the demographic information of species when information about the phylogeny is available, to perform phylogenetic comparative analyses (Revell, 2010; Revell, 2012), the final dataset included 259 species of Crocodilia (23) and Testudines (236) with this resource

Demographic data of reptiles are scarce and contain missing values (Conde et al. 2019; Fig S2). As principal component analyses require full data (*i*.*e*. non-gappy data; Rubin 1976, Nakagawa 2004), we carried out phylogenetic imputations to fill in the gaps in the life history traits for the studied species (see Appendix S2. Missing data). To impute the missing life history traits in our dataset, we used the R package mice (Van Buuren & Groothuis-Oudshoorn, 2011), which uses multiple imputation and the addon-phylomice to include phylogenetic information. This approach has been shown recently to correctly impute missing data in similar datasets better than other available approaches (Murray, 2018). Briefly, this approach uses Fully Conditional Specification (FCS) of the imputation model, which specifies the multivariate imputation model on a variable-by-variable basis by a set of conditional densities, one for each incomplete variable (Van Buuren & Groothuis-Oudshoorn, 2011). As suggested by previous studies (Janssen et al., 2010), we included three more traits in the imputation analyses in order to have more robust estimations of the imputed data (we included body weight and body size, which is available for all the species, and incubation time available for 16% of the species from Amniote Database). Such as there are some stochasticity in the imputation method, we then created 40 imputed datasets (the percentage of gaps in the data were 38%) and ran analyses on each separately. Like this approach used the phylogenetic information, we evaluated the phylogenetic signal. We first estimated the phylogenetic signal for each trait separately using Pagel’s *λ*, which describes the strength of phylogenetic relationships on trait evolution under a Brownian motion model (Freckleton 2000). Pagel’s *λ* ranges between 0, when the patterns in the traits cannot be explained by the employed phylogeny, and 1 when the observed patterns in traits are tightly correlated with the placement of species in the phylogeny. Most of the selected traits showed a strong phylogenetic signal (> 0.8; see Table S3), so we used the imputed values for traits in further analyses. We evaluated the similarity between imputed and non-imputed data using density plots for each trait. Moreover, we analysed the position in the functional spectra using Procrustres analyses between imputed data and a subset where a maximum of two traits were imputed (see Appendix S2. Imputation validation).

### Exploring the relation between the axes of life history strategies

To identify potential differences between the patterns of association among life history traits for Testudines and Crocodilia species while simultaneously assessing non-independence of lineages, we used a phylogenetically informed PCA (*p*PCA; Revell, 2009). *p*PCA is a multivariate analysis that reduces the number of variables of interest due to their plausible correlation. The *p*PCA considers the correlation matrix of species’ traits while accounting for phylogenetic relationships and estimate Pagel’s *λ*. The *p*PCA was estimated using the R package *phytools* (Revell, 2012), assuming a Brownian motion model of evolution (Revell, 2010). Life history trait data were log□transformed to fulfil normality assumptions of PCA and z□transformed to mean = 0, and SD = 1 (Legendre & Legendre, 2021). We used the Kaiser criterion (Kaiser, 1960) after optimisation through varimax rotation to determine the number of axes necessary to explain a substantial amount of variation. Namely, the axes we retained had an associated eigenvalue >1.

As body weight is strongly correlated with life history traits (Gaillard et al., 1989; Bielby et al. 2007; Capdevila et al., 2020), we accounted for this potential effect in our multivariate analyses too. Out of the multiple possible ways to account for body size or weight in life history analyses, we used the residuals of phylogenetic regression between body size and each trait (Revell, 2009). We collected most of adult body mass (g) data from the studies by Myhrvold et al. (2015) and Colston et al. (2020), moreover, we collected the remained data from previous literature for all our 259 species.3

### Estimating the functional spectra

To describe the probabilistic distribution of the species within the functional spaces, we used the main axes of functional trait variation by *p*PCA corrected by size (*i*.*e*., residuals of regressions). We performed multivariate kernel density using the ‘*TPD’* and ‘*ks’* R packages (Duong, 2007, 2014; Carmona, 2019; Carmona et al., 2019) to estimate the functional spectra. The kernel was a multivariate normal distribution for each species centered in the location of the species in *p*PCA and bandwidth chosen using unconstrained bandwidth selectors from the ‘*Hpi’* function in the ‘*ks’* package (Duong & Hazelton, 2003). The grouped kernels for all species drive into the continuous TPD function (Carmona et al., 2017; 2019). According to Carmona et al. (2021) and Toussaint et al. (2021), we divided the continuous functional space into a two-dimensional grid composed of 200 equal sized cells per dimension. We estimated the value of the TPD function for the 40,000 cells. The value of the TPD function represent the density of species in that particular region of the functional space (i.e., species with similar life history strategy). Results of TPD were represented graphically with the contours containing 50, 60, 70, 80, 90, and 99% of the total density of species.

### Threats to the studied species

To identify causes of recent declines in Testudines and Crocodilia species we reviewed published information. The main threats described for these species were i) habitat loss, fragmentation, and degradation (Luiselli, 2009), ii) over-collection of individuals and their eggs for food consumption (Gong et al., 2017), iii) unsustainable or illegal international trade, as well as over-collection for the trade in medicines (Sung & Fong, 2018), iv) climate change (Gibbons et al., 2000); v) emerging diseases (Jacobson et al., 2014; Tompkins et al., 2015). We collected the information on the threats to each species from Stanford et al. (2020), Bonin et al. (2006) and IUCN Red List (IUCN, 2020). We did dichotomous variables when one of these threats were mentioned. We were not able to quantify the effect of the threat, because effect size is only available for the Critically Endangered species of Stanford et al., (2020). To convert the threat descriptions of the Red List into our broader categories, we used the following process: for habitat disturbances we considered the Red List threat classifications “residential & commercial development”, “agriculture & aquaculture” and “natural system modifications”; for climate change the classification “climate change and extreme weather”; and for disease the classification “invasive and other problematic species, genes & diseases”.

### Effects of extinctions of threatened species on global functional diversity

To identify the loss of functional diversity by threatened species extinctions, we simulated the disappearance of species according to their Red List category (IUCN, 2020). The species classified among the threatened species categories (i.e. Critically Endangered (CR), Endangered (EN) and Vulnerable (VU) have a higher risk of extinction than the rest of species: Near Threatened (NT), Least Concern (LC); the extinction risk of Data Deficient (DD) species is unknown. In our analysis, we removed threatened species and estimated the resulting shifts in functional diversity (Toussaint et al., 2021). Firstly, we removed the species with a higher risk of extinction (-CR), then we removed successively the species with lower risks (-EN includes the extinction of species classified as a CR and EN, -VU includes CR, EN and VU, and -NT includes CR, EN, VU and NT). These extinction scenarios represented a gradient of extinction risk from the persistence of all species to a more dramatic scenario where all threatened species (including the NT species) went extinct. We compared the TPD function considering all the species assessed by IUCN (current spectra of functional diversity), and the TPD function after removing the species in the different scenarios. We can compare TPD functions because they are probability density functions, which means that they integrate to 1 across the whole functional space, regardless of the number of species considered (Carmona et al. 2016). To reduce the potential effect of outliers in the functional space, we applied a quantile threshold of 99%. We quantify how much of the functional spectra is lost after the extinction scenarios by estimating which functional space cells become empty after extinctions. The difference in the amount of space occupied before and after extinctions (TPD function of the current spectra – TPD function after removing the threatened species) represented the loss of functional diversity for Testudines and Crocodilia. We calculated this loss in the functional space for all the extinction scenarios (-CR, -EN, -VU, and -NT).

To assess if the impacts of each extinction scenario were different from what would be expected if extinction risk is not related to life history traits, we also compared the observed changes in functional diversity to a null model where the extinct species were randomly selected within the pool of species. For each scenario of extinction risk, we compared the functional diversity to 999 losses of functional diversity where the same number of threatened species were randomly selected among the pool of species. This strategy allowed us to understand whether the extinction in the different scenarios reduces more or less than expected the functional diversity. For each scenario, we created 999 TPD functions simulating cases in which the same number of species were lost at random from the total set of IUCN-assessed species. We compared the 5% and 95% percentile of the random simulation with the value of loss of functional diversity per each extinction scenario to estimate if the values were significantly different. In this case, higher than expected reductions in functional diversity would mean that the species that are going extinct in the considered scenario are relatively unique in terms of their functional traits, whereas lower than expected reductions would imply that the species going extinct are mostly functionally redundant.

After examining the loss of functional diversity, we plotted the extinction risk of species within the functional spaces, using the species assessed by IUCN. Using binomial smoother-based GAM (Wood, 2006) with the R package *mgcv* (Wood, 2011), we analysed the relationship between extinction risk (1: high-risk and 0: nonthreatened or low-risk) and the position in the functional space (PC axes). We then represented the predictions of the models (including the 95% confidence intervals of the means) to visually examine how life history strategies affect the probability of species being threatened (Carmona et al., 2021).

### Effects of anthropic threats on global functional diversity

To identify the effect of the different threats in the functional spectra, we simulated scenarios where extinctions were based upon species reported as affected by specific threats. We did two comparisons: 1) the TPD function considering all the species (current spectra of functional diversity) and TPD function after removing the species affected by specific threat (habitat disturbance, trade, local consumption, climate change or disease) and, 2) the TPD function considering all the IUCN-assessed species, and the TPD function after removing the threatened species (CR, EN, or VU) affected by each specific threat. Using a comparison between TPD functions similar to the one explained above, we estimated the potential loss of functional diversity attributable to each threat and evaluated the differences between them. We also compared the observed changes in functional diversity using a null model to assess if the impacts of each extinction scenario were different from what would be expected by chance. For each threat, we compared the loss of functional diversity with the 999 losses of functional diversity where the same number of species were randomly selected among the pool the species. We did these comparisons both considering the extinction of all the species affected by each threat and considering only the extinction of threatened species.

To visualise the relationship between life history strategies and threats, we mapped the probability of a species being affected by a specific threat within the functional spaces. We used binomial smoother-based GAM (Wood, 2006) with the R package *mgcv* (Wood, 2011) to identify the relationship between being affected by each threat (1: affected and 0: nonaffected) and the position in the functional space (PCA axes). We represented the predictions of the models (including the 95% confidence intervals of the means) when the relation between threat and life history strategy was significant.

## ACKNOWLEDGEMENTS

RCRC was supported by a post□doctoral grant funded by the Regional Valencian Government (APOSTD/2020/090) hosted by RSG. RSG was supported by a NERC IRF grant (NE/M018458/1). RCRC, EG, and AG were supported by the Spanish Ministry of Science and European Regional Development Fund (PID2019-105682RA-I00/AEI/10.13039/501100011033). CPC was supported by the Estonian Research Council (PSG293). Illustrations by Carmen Cañizares (@canitailustradora)

