## Supplementary Information for "Anthropogenic impacts on threatened species erode functional diversity in turtles and crocodilians"

### The extinction of threatened species by anthropogenic threats endangers the functional diversity of turtles and crocodilians

Rodríguez-Caro, R.C., Graciá, E., Blomberg, S. P., Cayuela, H., Grace, M., Carmona, C.P., Pérez-Mendoza, H.A., Giménez, A., Salguero-Gómez, R.

#### Index

**Appendix S1. Extended results** **P. 2**

Table S1 P. 2

Figure S1 P. 3

**Appendix S2. Extended methods** **P. 4**

Data selection P. 4

Missing data P. 9

Imputation validation P. 11

Body mass data P. 15

Phylogenetic signal P. 22

**References** **P. 23**

#### Appendix S1: Extended results

**Table S1. Loadings of phylogenetically-corrected principal component analysis (pPCA), also corrected by body mass, for the 236 species of Testudines and 23 species of Crocodilia examined in this study.** Only the first two PC axes are shown, as only they raised associated eigenvalues >1, indicating that they explain sufficient observed variation in life history traits (Legendre & Legendre, 2012). Bold numbers indicate loading absolute values >0.50. Outputs correspond to the mean values of 40 imputed data sets. Pagel's $\lambda$ indicates the extent to which patterns are explained (=1) or not (=0) by phylogenetic relationships. Variance estimates (±) correspond to standard deviation.

| Life history trait | Symbol | PC1 | PC2 |
|---|---|---|---|
| Adult survival | *Sa* | 0.03 ± 0.08 | 0.01 ± 0.12 |
| Juvenile survival | *Sj* | 0.04 ± 0.17 | -0.06 ± 0.33 |
| Age at sexual maturity | *Lα* | 0.03 ± 0.09 | -0.03 ± 0.10 |
| Clutch size | *CS* | **0.56 ± 0.07** | **-0.767** ± 0.08 |
| Mean number of clutches per year | *CN* | -0.13 ± 0.08 | 0.191 ± 0.14 |
| Maximum lifespan | *ML* | **0.92 ± 0.04** | 0.377 ± 0.10 |
| **Proportion of variance explained** | | 39.1% ± 1.87% | 23.1% ± 1.20% |
| **Cumulative proportion of variance explained** | | 39.1% | 62.2% |
| **Pagel's $\lambda$** | | 0.674 ± 0.030 | |

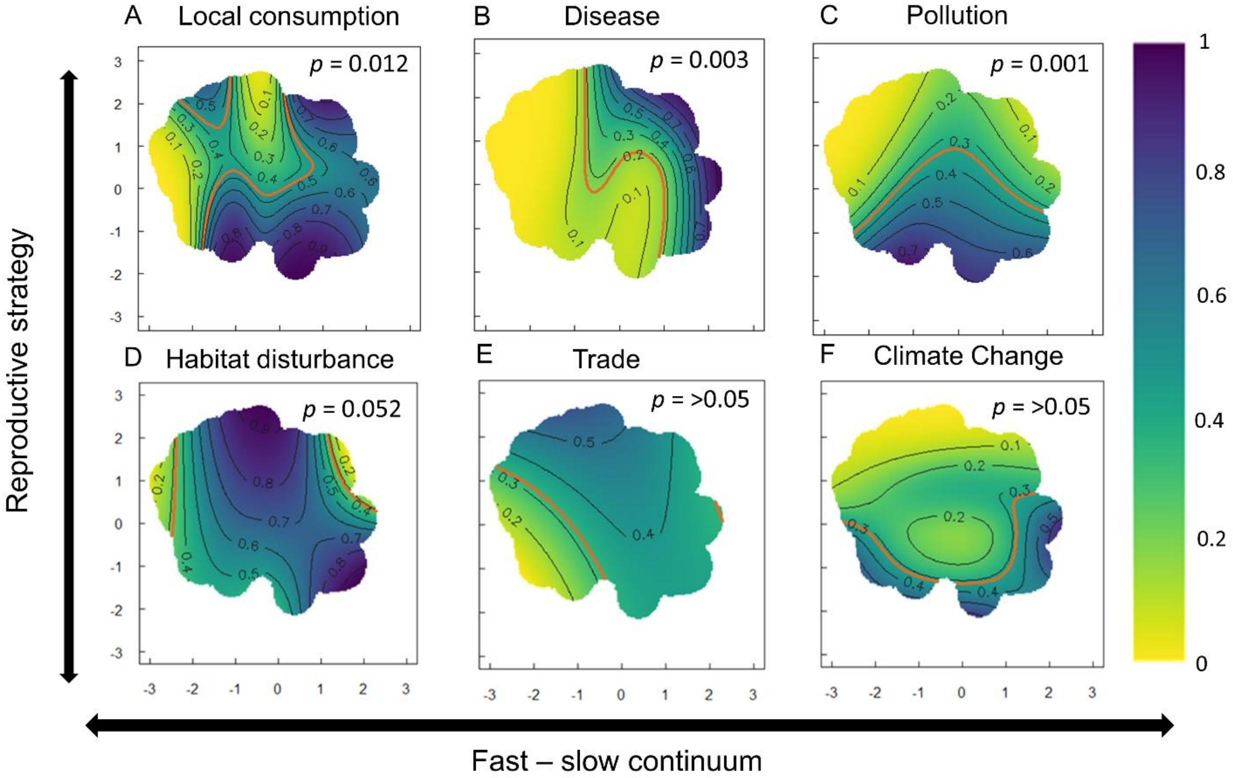


**Fig S1. The life history strategies of Testudines and Crocodilia predict the vulnerability to extinction due to some but not all considered threats.** Probability of species being affected by (A) local consumption, (B) disease, (C) pollution, (D) habitat disturbances, (E) unsustainable global trade, and (F) climate change, according to GAMs (with binomial distribution) using the position of species in the two-dimensional functional space as predictors. Local consumption, disease, and pollution can predict the extinction of some life history strategies. Other risks, such as climate change, global trade, or habitat disturbance, do not show a significant relation with the species' life history strategie**s.** Yellow tones indicate lower risk of extinction due to each threat, whereas purple tones indicate higher risk. P values associated with each GAM are shown at the top-right corner of each panel. The red contour lines indicate the average threat probability. We consider only species for which threats are known (n = 251 species).

### Appendix S2: Extended methods

#### *Data Selection*

We drew life history data from the COMADRE Animal Matrix Database (Salguero-Gómez et al. 2016), DATLife Database (2021), Amniote Life History Database (Myhrvold et al. 2015), and the reviews by Allen et al. (2017), Pfaller et al. (2018) and Reinke et al. (under review).

COMADRE is an open-access database that includes matrix population models (MPMs, Caswell, 2001) of 414 animal species around the globe (Salguero-Gómez et al., 2016). MPMs describe the lifecycle transitions of individuals across discrete state classes (*i.e.*, age/stage/size classes). We selected MPMs of species in the orders Testudines and Crocodilia for which the MPMs: (i) were parameterised with field data from natural populations, *i.e.* undisturbed, unmanipulated populations, so their life history traits represent those of natural populations; (ii) encompass the entire life cycle of the species (*i.e.* include rates of survival, development, and reproduction along the full life cycle), so we could then calculate life history traits that require full life cycle information; (iii) have three or more life cycle stages, since lower-dimension MPMs lack the necessary resolution to estimate life history traits (Salguero-Gómez & Plotkin, 2010); (iv) in the case of more than one study for the same species, the MPM with overall greater temporal replication was determined to better represent the species' demography. MPMs in COMADRE contain three sub-matrices necessary to quantify life history traits (Caswell 2001): a state-specific survival sub-matrix ($\boldsymbol{U}$), a stage-specific per capita reproduction sub-matrix ($\boldsymbol{F}$), and a stage-specific per capita clonality sub-matrix ($\boldsymbol{C}$). The stage-classified population projection matrix is then defined as $\boldsymbol{A} = \boldsymbol{U} + \boldsymbol{F} + \boldsymbol{C}$. In our study species, $\boldsymbol{C} = (0)_{n\times n}$ (where $n$ = life cycle stage number), as these taxonomic groups do not reproduce clonally. This set of criteria resulted in 24 MPMs for five Crocodilia species and 19 Testudines.

DATLife (Demography Across the Tree of Life) is an open-access database that includes life tables, longevity, and age at maturity for over 5,500 species (DATLife Database, 2021). The associated life tables contain the age-specific survivorship ($l_x$) and age-specific fertility ($m_x$) in populations where individual age ($x$) is known. We selected the life tables of species in the order Testudines and Crocodilia that were parameterised strictly with field data from wild populations and, as for the COMADRE data, in the case of more than one study available for the same species, we selected the study with the longest temporal replication. We obtained life history trait data for 122 species of Crocodilia (14) and Testudines (108) from this resource.

The Amniote Life History Database (Amniote) includes life history information of 21,223 species of birds, mammals, and reptiles (Myhrvold et al. 2015). Amniote pulls information from empirical peer-reviewed studies on individual species, macroecological studies of multiple species, existing life history databases, published books, and other compilations. From Amniote, we obtained primarily reproductive life history traits (*e.g.*, age at maturity for males and females, clutch size, clutches per year, duration of gestation, duration of weaning, incubation time, egg mass, inter-clutch interval), survival-related life history traits (*e.g.*, longevity and maximum lifespan), as well as biometric information (*e.g.,* body mass, hatching weight, male and female body mass, straight ventral length for males, females and hatchings). Where multiple values are available for a species, Amniote reports the median value (Myhrvold et al. 2015). We obtained data from 237 species of Crocodilia (23) and Testudines (214) from this resource.

To complement the information about life history traits obtained from the aforementioned databases, we carried out a search of peer-reviewed studies that present demographic data for tortoises, turtles, crocodilians and alligators. Allen et al. (2017), Pfaller et al. (2018) and Reinke et al. (under review) review and compile information about reproduction and survival of reptiles into open access databases. Allen et al. (2017) collected life history data about reproduction and biometric information by combining existing life history databases and supplementing these with additional data from the primary literature of 5,716 amphibian and 9,046 reptile species. In cases where Allen et al. (2017) and the Amniote database reported different values for a species' traits, we selected the information of Allen et al. (2017) for our analysis because it the data were more recent. Pfaller et al. (2018) published a review compiling annual survival probability estimates through capture-recapture data for adult sea turtles. Reinke et al. (under review) collected capture-recapture information to develop survival estimates for 45 reptile species.

To calculate the life history strategies of Testudines and Crocodilia, we collected/estimated the life history traits according to the different datasets:

- Mean number of clutches per year and clutch size (*CN* and *CS*, respectively) were obtained from Amniote (Myhrvold et al. 2015) and Allen et al. (2017). Allen et al. (2017) includes several data (including Amniote) and, when they found multiple records of the same trait for a species, the average of the species' trait was estimated by taking the mean of unique records per species. When the same species was present in both datasets, we selected the value in Allen et al. (2017).
- Adult and juvenile survival (*Sa* and *Sj*, respectively) were calculated from the MPMs in COMADRE (Salguero-Gómez et al., 2016a), life tables in DATLife (2021), and from direct estimates of capture-recapture published studies (such as Pfaller et al., 2018; Cayuela et al. 2019). Adult survival (*Sa*) from MPMs was estimated as the column sum of the matrix ***U*** of the stages representing reproductive individuals represented in the sub-matrix ***F*** (considering adults, all ages/stages after the first reproduction). We used arithmetic averages for the resulting stage-specific

survival values, rather than weighting them by the stable stage distribution (*e.g.*, Roper et al., 2021) because we collected juvenile and adult data from other sources where calculating this weighted mean would not feasible, thus rendering comparisons impossible. In DATLife, we used age-specific fertility to determine the reproductive ages, and then estimated adult survival as the arithmetic average of age-specific survival rates after the first reproduction. We estimated juvenile survival (*Sj*) using a similar approach: as the average of the sum of the columns that represent juveniles (all the stages prior to the first reproduction) in the pertinent sub-matrix ***U*** in COMADRE. In DATLife, we estimated the average of non-reproductive age- or stage-specific survival (prior to the first reproduction) of the life tables.

- Age at maturity ($L\alpha$) and maximum lifespan (*ML*) were estimated with the MPMs from COMADRE or obtained directly from the databases with this information available (such as Amniote or DATLife). In COMADRE, $L\alpha$ was calculated using age-from-stage decompositions (Caswell 2001, p. 124). Briefly, we defined the reproductive stages as those columns of the ***F*** sub-matrix that contain values greater than zero. We estimated the mean time between birth and the first entry into the reproductive stage. This conditional mean is obtained by creating an absorbing state corresponding to the event of reproducing at least once before death, with a Markov chain, and calculating the mean time to absorption (Caswell, 2001). *ML* was estimated from the MPMs of COMADRE projecting 100 individuals in the first state and iterating up to 1000 years to identify the first year with fewer than one individual in the virtual cohort. We also used the databases Amniote and DATLife because they contain information about $L\alpha$ and *ML* of most of the examined species. When several values from different datasets for one species and life history trait were available, we selected the lowest value for $L\alpha$ and the highest values for $ML$. The rationale behind this choice is that the lowest value of $L\alpha$ identifies the most likely first age at maturity reported in the life cycle of this species, whereas the highest values of $ML$ approximate the maximum values of this trait reported for the species of interest in wildlife populations.

*Missing data*

For some species, specific life history traits were not directly available from databases/literature, nor were we able to calculate them with the demographic data that were available. This resulted in gaps in our dataset of species' life history traits. Thus, we explored the patterns of missing values for Testudines and Crocodilia species (Fig. S2). The traits with the highest proportion of missing values were juvenile survival (12.7% of all species) and adult survival (13.5%). Clutch number was available for 37.8% and age at maturity for 44%. The traits best represented in the database were maximum lifespan (79.9%) and clutch size (97.7%). Overall, the percentage of gaps was 38%, then we imputed around 1 dataset for every percent missing, so we imputed 40 datasets.

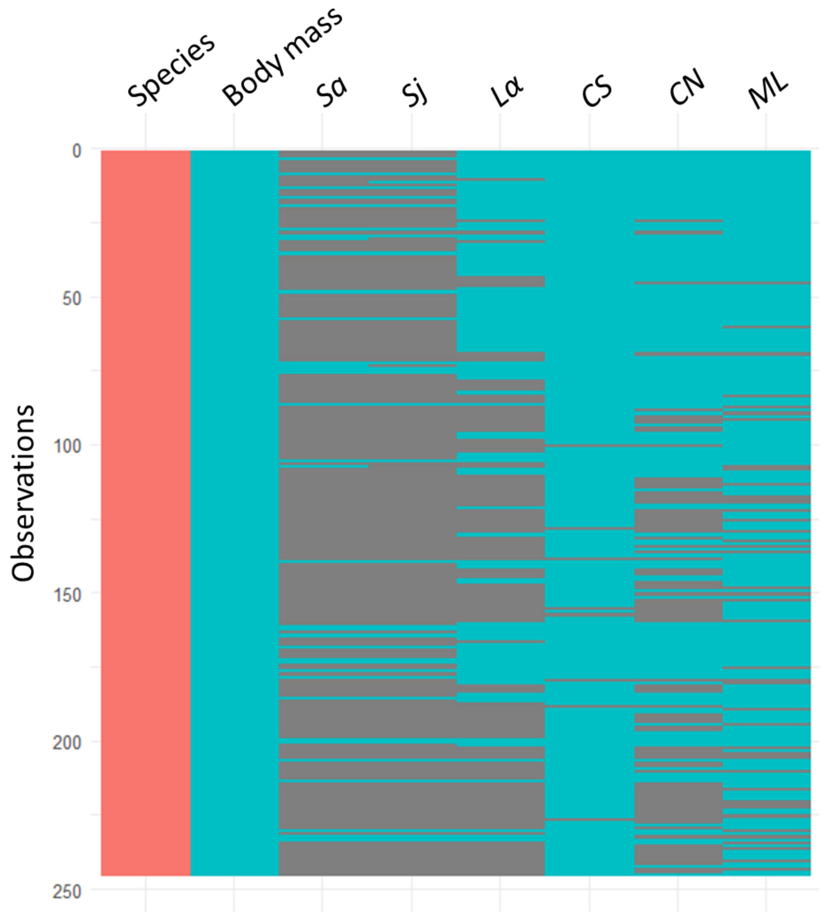

**Fig S2. Proportion of missing life history trait values for the studied species**. Despite our search of literature and existing datasets, we were unable to find values for all life history traits for all species ($N$ = 256). The life history traits are: adult survival (Sa), juvenile survival (Sj), maximum lifespan (ML), age at sexual maturity (Lα), mean of number of clutches per year (CN), clutch size (CS). Red values are character records such as the name of the species, blue values are numeric information and grey values are gaps of information (NA data).

*Imputation validation*

Because the *p*PCA analyses require complete datasets, we imputed the missing data using the add-on *phylomice* to *Mice* package (Van Buuren & Groothuis-Oudshoorn, 2011), as detailed in the Methods section *Phylogeny and data imputation* of our main manuscript.

We compared the distribution of the imputed data versus original data (with no missing values) using density plots (Fig S3). Visually we can identify that the data from the 40 imputed datasets matched with the values with the original data.

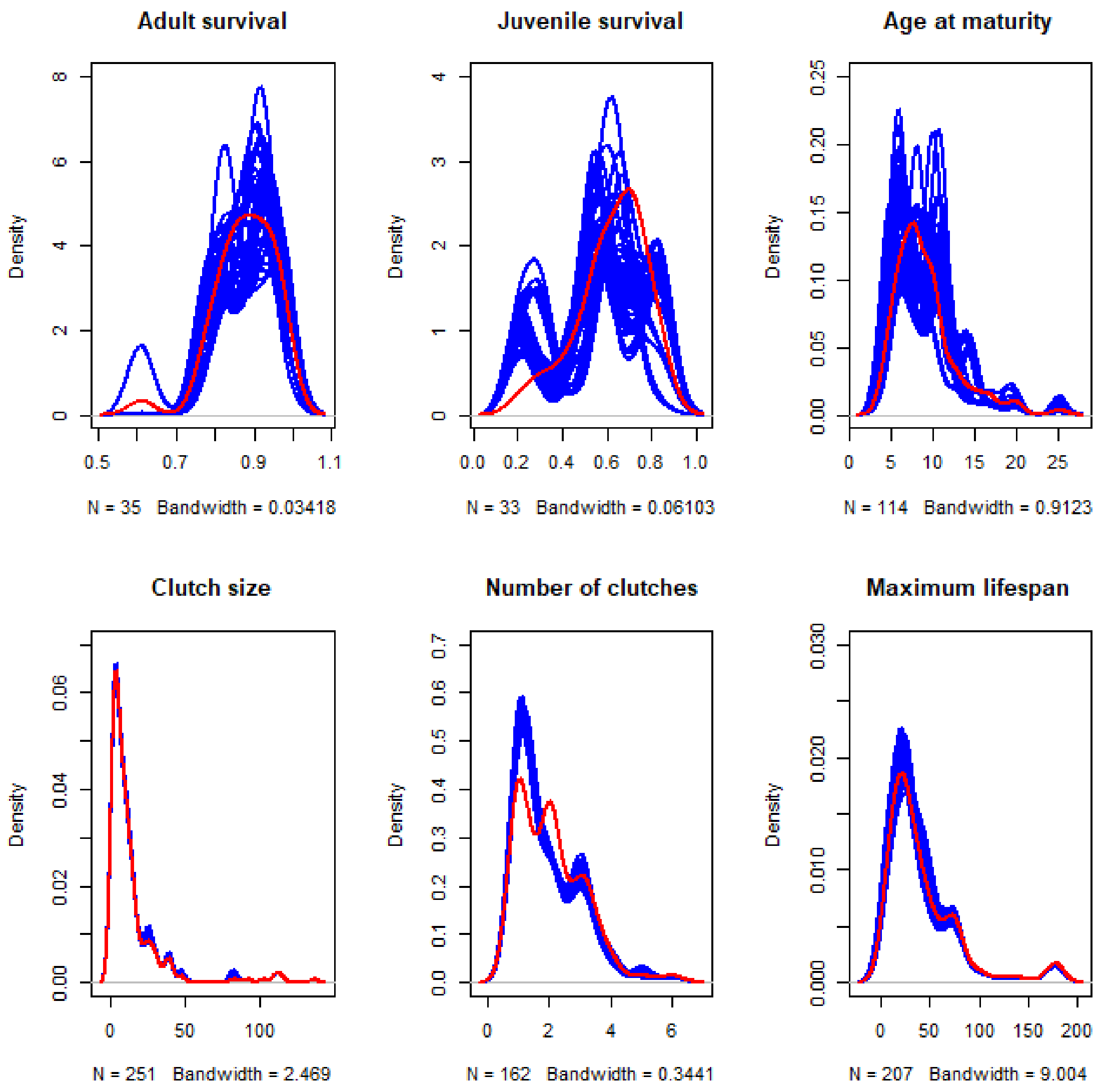

**Fig. S3. Comparison density plots between imputed versus non-imputed data for each life history trait.** In red, the density of values of observed data, in blue the datasets estimated in each imputation (including observed and imputed values). N is the number of observed values per trait, for example, in adult survival we have information available for 35 species and 251 species for clutch size. Bandwidth was fixed according to the observed data in the density plot, in other words, the bandwidth used in the red line (observed data) was the same that it was used for all imputations.

To evaluate the performance of the imputation method utilised in our study, we compared the main results of the *p*PCA with imputed data versus a subset of the dataset where a maximum of two traits were imputed. This new subset encompassed 108 species and we have no gaps for *CS* and *Lα,* the percentage of gap information for *CN* was 0.9 %, for *ML* was 1.9%. However, the data with the highest empty values are those related to survival (*Sa* = 68.5% and *Sj* = 69.4%. ). We performed a *p*PCA using this subset of 108 species. Visually we could see that the main trends of the analyses are similar (Fig. S4). The Pagel's $\lambda$ of the new model with few imputed data was 0.825, slightly higher than the model with imputed data. The proportion of variance explained by the *pPCA* was also higher for the first axis, PC1 = 42.39%, and it was similar for the second axis, PC2 = 25.28%. This similarity in the results support the use of the imputed data.

a)

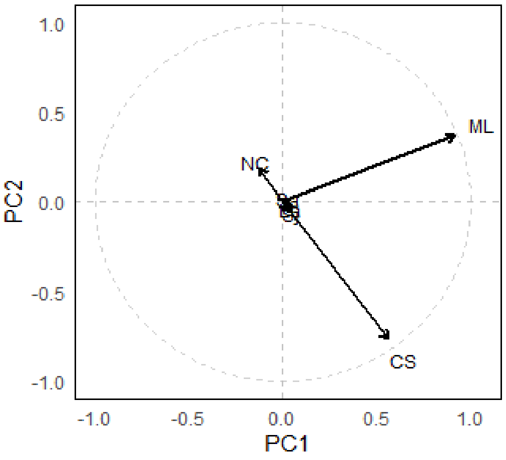


b)

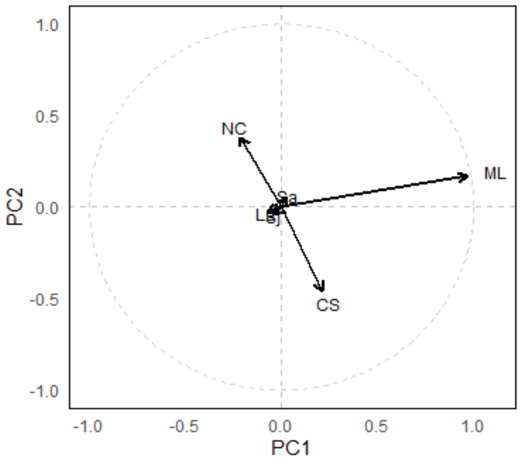

**Fig S4. Similar loadings of *pPCA* corrected by size are shown in the two models: a) using imputed data to fill gaps, so that all species could be included (N = 259); and b) using only species with a maximum of two imputed traits (N = 108).** Representation of the axis of variation for the model used in the manuscript (a) and the model with a selection of species without imputed data (b).

To quantify the differences between both *p*PCA analyses, we did a Procrustes analysis. This analysis shows up to what point the position of the common species (108 species) remains constant between the two spaces. To carry on these analyses, we used the function procrustes from the *vegan* package in R (Oksanen et al., 2013). The results showed high correlation in a symmetric Procrustes rotation (0.9656) between both datasets and the sum of squared distances between paired points in the ordination space was 0.067 (Fig. S5). The permutational test of the significance of the Procrustes was done with 9999 permutations, and showed significant relations between the scores of imputed database and scores where a maximum of two traits were imputed ($p < 0.001$). This result means high correlation between both *p*PCA analyses, supporting the use of the imputed data in the analyses.

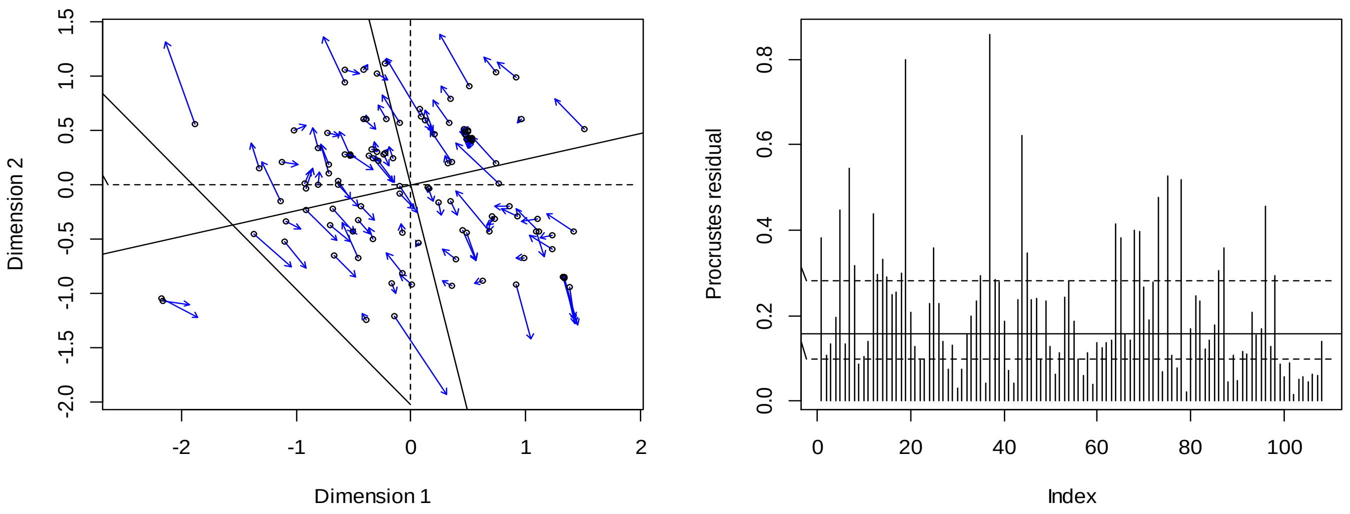

**Fig S5. Representation of the Procrustes errors.** On the left, visual indication of the degree of match between the two spaces. Symbols show the position of the samples in the *p*PCA scores with a maximum of two traits imputed per species, and arrows point to their positions according to the score of the *p*PCA with the imputed data. The plot also shows the axis rotation between the two ordinations. On the right, the plot shows the residuals for each species. The horizontal lines, from bottom to top, are the 25% (dashed), 50% (solid), and 75% (dashed) quantiles of the residuals.

#### *Body mass data*

Body mass is highly correlated with many life history traits (Gaillard et al. 1989; Healy et al. 2019), and therefore in this manuscript our multivariate analyses account for body mass. We collected the female body mass values for all studied species using the datasets described in the section *Data selection* of the Supplementary Information. We also searched for body mass data in the literature, including data presented under previously taxonomically accepted names for a given species. When we were not able to find a value for body mass, we estimated body mass using the total length of the species (using the information of the same clade, *i.e.* family or genus). We conducted simple regression between body mass and total length because these traits are highly correlated, we used all the species corresponding to the taxonomic genus or family, according to the number of species available in each taxonomic group. We estimated body mass using this approach for 10 of the 259 species included in the analyses (Table S2).

**Table S2. List of species used in the manuscript for which females body mass (g.) was available.**

| Order | Family | Species | Body Mass | Database |
|---|---|---|---|---|
| Crocodilia | Alligatoridae | *Alligator mississippiensis* | 47800 | Amniote Database |
| | | *Alligator sinensis* | 10750 | Amniote Database |
| | | *Caiman crocodilus* | 10550 | Amniote Database |
| | | *Caiman latirostris* | 14600 | Amniote Database |
| | | *Caiman yacare* | 10550 | Amniote Database |
| | | *Melanosuchus niger* | 82000 | Amniote Database |
| | | *Paleosuchus palpebrosus* | 5900 | Amniote Database |
| | | *Paleosuchus trigonatus* | 7500 | Amniote Database |
| | Crocodylidae | *Crocodylus acutus* | 76700 | Amniote Database |
| | | *Crocodylus intermedius* | 107900 | Amniote Database |
| | | *Crocodylus johnsoni* | 19500 | Amniote Database |
| | | *Crocodylus mindorensis* | 38400 | Amniote Database |
| | | *Crocodylus moreletii* | 31700 | Amniote Database |
| | | *Crocodylus niloticus* | 94200 | Amniote Database |
| | | *Crocodylus novaeguineae* | 39900 | Amniote Database |

| | | | | |
|---|---|---|---|---|
| | | *Crocodylus palustris* | 42700 | Amniote Database |
| | | *Crocodylus porosus* | 78700 | Amniote Database |
| | | *Crocodylus rhombifer* | 57500 | Amniote Database |
| | | *Crocodylus siamensis* | 42500 | Amniote Database |
| | | *Mecistops cataphractus* | 50500 | Amniote Database |
| | | *Osteolaemus tetraspis* | 18800 | Amniote Database |
| | | *Tomistoma schlegelii* | 119000 | Amniote Database |
| | Gavialidae | *Gavialis gangeticus* | 147000 | Allen et al. 2017 |
| Testudines | Carettochelyidae | *Carettochelys insculpta* | 10050 | Amniote Database |
| | Chelidae | *Acanthochelys pallidipectoris* | 369.2 | Amniote Database |
| | | *Acanthochelys spixii* | 330.87 | Fraxe Neto et al. 2011 |
| | | *Chelodina longicollis* | 1293 | Amniote Database |
| | | *Chelodina mccordi* | 2340 | Colston et al. 2020 |
| | | *Chelodina novaeguineae* | 1006 | Amniote Database |
| | | *Chelodina reimanni* | 1006 | Amniote Database |
| | | *Elseya dentata* | 4295 | Amniote Database |
| | | *Emydura macquarii* | 1740 | Amniote Database |
| | | *Emydura subglobosa* | 1030 | Amniote Database |
| | | *Emydura tanybaraga* | 1030 | Amniote Database |
| | | *Hydromedusa maximiliani* | 325 | Regis and Meik 2017 |
| | | *Hydromedusa tectifera* | 1155 | Regis and Meik 2017 |
| | | *Mesoclemmys dahli* | 777 | Amniote Database |
| | | *Mesoclemmys gibba* | 706.8 | Amniote Database |
| | | *Mesoclemmys nasuta* | 2180 | Cunha et al 2019 |
| | | *Mesoclemmys tuberculata* | 657.4 | Regis and Meik 2017 |
| | | *Mesoclemmys vanderhaegei* | 453.29 | Regis and Meik 2017 |
| | | *Mesoclemmys zuliae* | 1983 | Amniote Database |
| | | *Myuchelys bellii* | 1628 | Fielder et al 2015 |
| | | *Myuchelys georgesi* | 977.3 | Regis and Meik 2017 |
| | | *Myuchelys latisternum* | 1377 | Amniote Database |
| | | *Phrynops geoffroanus* | 2233.3 | Regis and Meik 2017 |
| | | *Phrynops hilarii* | 2380 | Amniote Database |
| | | *Phrynops williamsi* | 4740.97 | Encyclopedia of Life |
| | | *Platemys platycephala* | 325 | Amniote Database |
| | | *Pseudemydura umbrina* | 324.1 | Amniote Database |
| | | *Rheodytes leukops* | 1691 | Amniote Database |
| | | *Rhinemys rufipes* | 1265.96 | Estimated |
| | Cheloniidae | *Caretta caretta* | 92400 | Amniote Database |
| | | *Chelonia mydas* | 128100 | Amniote Database |
| | | *Eretmochelys imbricata* | 62300 | Amniote Database |
| | | *Lepidochelys kempii* | 92400 | Amniote Database |
| | | *Lepidochelys olivacea* | 116300 | Amniote Database |
| | | *Chelydra rossignonii* | 3875 | Amniote Database |
| | | *Chelydra serpentina* | 3875 | Amniote Database |
| | | *Macrochelys temminckii* | 23500 | Amniote Database |

| | | | |
|---|---|---|---|
| Dermatemydidae | *Dermatemys mawii* | 15660 | Amniote Database |
| Dermochelyidae | *Dermochelys coriacea* | 372500 | Amniote Database |
| Emydidae | *Chrysemys dorsalis* | 378 | Amniote Database |
| | *Chrysemys picta* | 378 | Amniote Database |
| | *Clemmys guttata* | 165.4 | Amniote Database |
| | *Deirochelys reticularia* | 910.5 | Amniote Database |
| | *Emys orbicularis* | 1096.49 | Allen et al. 2017 |
| | *Glyptemys insculpta* | 968.2 | Amniote Database |
| | *Glyptemys muhlenbergii* | 132.1 | Amniote Database |
| | *Graptemys barbouri* | 1256 | Amniote Database |
| | *Graptemys caglei* | 1085.91 | Estimated |
| | *Graptemys ernsti* | 1519 | Aprox.[a] |
| | *Graptemys flavimaculata* | 1130 | Amniote Database |
| | *Graptemys geographica* | 1138 | Amniote Database |
| | *Graptemys gibbonsi* | 2334.5 | Amniote Database |
| | *Graptemys nigrinoda* | 415.7 | Amniote Database |
| | *Graptemys oculifera* | 918.15 | Amniote Database |
| | *Graptemys ouachitensis* | 1306.5 | Amniote Database |
| | *Graptemys pearlensis* | 1530.26 | Estimated |
| | *Graptemys pseudogeographica* | 1477 | Amniote Database |
| | *Graptemys pulchra* | 1519 | Amniote Database |
| | *Graptemys versa* | 1477 | Amniote Database |
| | *Malaclemys terrapin* | 886 | Amniote Database |
| | *Pseudemys alabamensis* | 3238.5 | Amniote Database |
| | *Pseudemys concinna* | 2992 | Amniote Database |
| | *Pseudemys gorzugi* | 2992 | Amniote Database |
| | *Pseudemys nelsoni* | 3738.5 | Amniote Database |
| | *Pseudemys peninsularis* | 3065.5 | Amniote Database |
| | *Pseudemys rubriventris* | 3238.5 | Amniote Database |
| | *Pseudemys texana* | 2992 | Amniote Database |
| | *Terrapene carolina* | 372 | Amniote Database |
| | *Terrapene coahuila* | 259.8 | Amniote Database |
| | *Terrapene nelsoni* | 372 | Amniote Database |
| | *Terrapene ornata* | 391 | Amniote Database |
| | *Trachemys callirostris* | 1854 | Amniote Database |
| | *Trachemys decorata* | 2522 | Amniote Database |
| | *Trachemys decussata* | 630.1 | Amniote Database |
| | *Trachemys dorbigni* | 1854 | Amniote Database |
| | *Trachemys gaigeae* | 1854 | Amniote Database |
| | *Trachemys grayi* | 1813 | Amniote Database |
| | *Trachemys nebulosa* | 1854 | Amniote Database |
| | *Trachemys ornata* | 1813 | Amniote Database |
| | *Trachemys scripta* | 1813 | Amniote Database |
| | *Trachemys stejnegeri* | 2012.95 | Encyclopedia of Life |

| | | | |
|---|---|---|---|
| | *Trachemys taylori* | 1854 | Amniote Database |
| | *Trachemys terrapen* | 4364.33 | Encyclopedia of Life |
| | *Trachemys venusta* | 1813 | Amniote Database |
| | *Trachemys yaquia* | 1854 | Amniote Database |
| Geoemydidae | *Batagur baska* | 17900 | Amniote Database |
| | *Batagur borneoensis* | 16900 | Amniote Database |
| | *Batagur dhongoka* | 7270 | Amniote Database |
| | *Batagur kachuga* | 21100 | Amniote Database |
| | *Batagur trivittata* | 21611.55 | Encyclopedia of Life |
| | *Cuora amboinensis* | 1000 | Amniote Database |
| | *Cuora aurocapitata* | 530.85 | Estimated |
| | *Cuora bourreti* | 499 | Amniote Database |
| | *Cuora flavomarginata* | 499 | Amniote Database |
| | *Cuora galbinifrons* | 1200 | Colston et al. 2020 |
| | *Cuora mccordi* | 375 | Amniote Database |
| | *Cuora mouhotii* | 850 | Colston et al. 2020 |
| | *Cuora picturata* | 1100 | Colston et al. 2020 |
| | *Cuora trifasciata* | 3000 | Aprox.[b] |
| | *Cuora yunnanensis* | 850 | Amniote Database |
| | *Cyclemys dentata* | 1250 | Amniote Database |
| | *Geoclemys hamiltonii* | 6000 | Regis and Meik 2017 |
| | *Geoemyda japonica* | 196 | Amniote Database |
| | *Geoemyda spengleri* | 196 | Amniote Database |
| | *Hardella thurjii* | 10500 | Regis and Meik 2017 |
| | *Heosemys annandalii* | 4000 | Andersen et al. 2021 |
| | *Heosemys grandis* | 4600.78 | Regis and Meik 2017 |
| | *Heosemys spinosa* | 950 | Amniote Database |
| | *Leucocephalon yuwonoi* | 1152.27 | Regis and Meik 2017 |
| | *Malayemys subtrijuga* | 3535 | Allen et al. 2017 |
| | *Mauremys annamensis* | 1717 | McCormack et al. 2004 |
| | *Mauremys caspica* | 912.01 | Allen et al. 2017 |
| | *Mauremys japonica* | 494.2 | Amniote Database |
| | *Mauremys leprosa* | 608 | Amniote Database |
| | *Mauremys mutica* | 319.5 | Amniote Database |
| | *Mauremys nigricans* | 912 | Regis and Meik 2017 |
| | *Mauremys reevesii* | 858 | Amniote Database |
| | *Mauremys rivulata* | 257.5 | Regis and Meik 2017 |
| | *Mauremys sinensis* | 1241 | Amniote Database |
| | *Melanochelys tricarinata* | 1800 | Regis and Meik 2017 |
| | *Melanochelys trijuga* | 760 | Amniote Database |
| | *Notochelys platynota* | 4364.33 | Encyclopedia of Life |
| | *Orlitia borneensis* | 12400 | Amniote Database |
| | *Pangshura smithii* | 919 | Amniote Database |
| | *Pangshura sylhetensis* | 1150 | Regis and Meik 2017 |
| | *Pangshura tecta* | 1000 | Regis and Meik 2017 |

| | | | |
|---|---|---|---|
| | *Pangshura tentoria* | 1184 | Amniote Database |
| | *Rhinoclemmys annulata* | 638.64 | Estimated |
| | *Rhinoclemmys areolata* | 731.5 | Amniote Database |
| | *Rhinoclemmys diademata* | 1126 | Amniote Database |
| | *Rhinoclemmys funerea* | 946 | Amniote Database |
| | *Rhinoclemmys melanosterna* | 2700 | Amniote Database |
| | *Rhinoclemmys nasuta* | 1284 | Amniote Database |
| | *Rhinoclemmys pulcherrima* | 944.6 | Regis and Meik 2017 |
| | *Rhinoclemmys punctularia* | 2344 | Encyclopedia of Life |
| | *Rhinoclemmys rubida* | 287 | Butterfield et al. 2018 |
| | *Sacalia bealei* | 329.6 | Lin et al. 2018 |
| | *Sacalia quadriocellata* | 284.95 | Regis and Meik 2017 |
| | *Siebenrockiella crassicollis* | 940 | Amniote Database |
| | *Vijayachelys silvatica* | 230 | Amniote Database |
| Kinosternidae | *Claudius angustatus* | 200 | Amniote Database |
| | *Kinosternon alamosae* | 145 | Amniote Database |
| | *Kinosternon baurii* | 143 | Amniote Database |
| | *Kinosternon chimalhuaca* | 266 | López-Luna et al. 2018[c] |
| | *Kinosternon durangoense* | 271.3 | Amniote Database |
| | *Kinosternon flavescens* | 271.3 | Amniote Database |
| | *Kinosternon hirtipes* | 202.6 | Amniote Database |
| | *Kinosternon integrum* | 474.4 | Amniote Database |
| | *Kinosternon scorpioides* | 266 | Amniote Database |
| | *Kinosternon sonoriense* | 326 | Amniote Database |
| | *Kinosternon subrubrum* | 152.35 | Amniote Database |
| | *Staurotypus salvinii* | 900 | Amniote Database |
| | *Staurotypus triporcatus* | 4200 | Amniote Database |
| | *Sternotherus carinatus* | 248 | Amniote Database |
| | *Sternotherus depressus* | 144 | Amniote Database |
| | *Sternotherus minor* | 154.5 | Amniote Database |
| | *Sternotherus odoratus* | 137.9 | Amniote Database |
| Pelomedusidae | *Pelomedusa subrufa* | 2273 | Amniote Database |
| | *Pelusios adansonii* | 1620 | Regis and Meik 2017 |
| | *Pelusios bechuanicus* | 4740.97 | Encyclopedia of Life |
| | *Pelusios castaneus* | 368.5 | Rawski and Józefiak 2014 |
| | *Pelusios castanoides* | 800 | Amniote Database |
| | *Pelusios chapini* | 3515.1 | Estimated |
| | *Pelusios nanus* | 311.93 | Encyclopedia of Life |
| | *Pelusios niger* | 1510 | Akani et al. 2015 |
| | *Pelusios rhodesianus* | 900 | Amniote Database |
| | *Pelusios sinuatus* | 7000 | Regis and Meik 2017 |
| | *Pelusios subniger* | 1232.64 | Encyclopedia of Life |
| | *Pelusios upembae* | 1210.35 | Estimated |

| | | | |
|---|---|---|---|
| | *Pelusios williamsi* | 2246.59 | Encyclopedia of Life |
| Platysternidae | *Platysternon megacephalum* | 305.5 | Regis and Meik 2017 |
| Podocnemididae | *Erymnochelys madagascariensis* | 4900 | Regis and Meik 2017 |
| | *Podocnemis erythrocephala* | 1412 | Regis and Meik 2017 |
| | *Podocnemis expansa* | 25800 | Amniote Database |
| | *Podocnemis lewyana* | 9599 | Amniote Database |
| | *Podocnemis sextuberculata* | 25800 | Amniote Database |
| | *Podocnemis unifilis* | 6380 | Amniote Database |
| | *Podocnemis vogli* | 2013 | Amniote Database |
| Testudinidae | *Aldabrachelys gigantea* | 33000 | Amniote Database |
| | *Astrochelys radiata* | 7955 | Amniote Database |
| | *Astrochelys yniphora* | 8800 | Amniote Database |
| | *Chelonoidis becki* | 43360.3 | Estimated |
| | *Chelonoidis carbonarius* | 6087.5 | Regis and Meik 2017 |
| | *Chelonoidis chathamensis* | 29954.54 | Estimated |
| | *Chelonoidis chilensis* | 3181 | Amniote Database |
| | *Chelonoidis darwini* | 36238.49 | Estimated |
| | *Chelonoidis denticulatus* | 3675 | Regis and Meik 2017 |
| | *Chelonoidis duncanensis* | 22000 | Chiari et al. 2017 |
| | *Chelonoidis hoodensis* | 28000 | Chiari et al. 2017 |
| | *Chelonoidis porteri* | 72000 | Chiari et al. 2017 |
| | *Chelonoidis vicina* | 61000 | Chiari et al. 2017 |
| | *Chersina angulata* | 715 | Amniote Database |
| | *Geochelone elegans* | 2500 | Amniote Database |
| | *Geochelone platynota* | 2500 | Amniote Database |
| | *Gopherus agassizii* | 2443 | Amniote Database |
| | *Gopherus berlandieri* | 1769.5 | Amniote Database |
| | *Gopherus flavomarginatus* | 85000 | McDonald 2008 |
| | *Gopherus polyphemus* | 2784 | Amniote Database |
| | *Homopus areolatus* | 289.4 | Amniote Database |
| | *Homopus femoralis* | 599 | Amniote Database |
| | *Indotestudo elongata* | 255 | Amniote Database |
| | *Indotestudo forstenii* | 967.5 | Amniote Database |
| | *Indotestudo travancorica* | 255 | Amniote Database |
| | *Kinixys belliana* | 1202 | Amniote Database |
| | *Kinixys erosa* | 958.4 | Regis and Meik 2017 |
| | *Kinixys homeana* | 690.7 | Regis and Meik 2017 |
| | *Kinixys lobatsiana* | 1202 | Amniote Database |
| | *Kinixys natalensis* | 1202 | Amniote Database |
| | *Kinixys nogueyi* | 1202 | Amniote Database |
| | *Kinixys spekii* | 617 | Hailey and Coulson 1996 |
| | *Kinixys zombensis* | 1202 | Amniote Database |
| | *Malacochersus tornieri* | 400 | Amniote Database |

| | | | |
|---|---|---|---|
| | *Manouria emys* | 30000 | Bonin et al. 2006 |
| | *Manouria impressa* | 3200 | Amniote Database |
| | *Psammobates geometricus* | 366.8 | Amniote Database |
| | *Psammobates tentorius* | 423 | Amniote Database |
| | *Pyxis arachnoides* | 398.1 | Regis and Meik 2017 |
| | *Pyxis planicauda* | 420 | Regis and Meik 2017 |
| | *Stigmochelys pardalis* | 20000 | Amniote Database |
| | *Testudo graeca* | 1430 | Amniote Database |
| | *Testudo kleinmanni* | 295 | Regis and Meik 2017 |
| | *Testudo marginata* | 2080 | Amniote Database |
| Trionychidae | *Amyda cartilaginea* | 2500 | Andersen et al. 2021 |
| | *Apalone ferox* | 20000 | Amniote Database |
| | *Apalone mutica* | 819 | Amniote Database |
| | *Apalone spinifera* | 4765 | Amniote Database |
| | *Chitra indica* | 108000 | Amniote Database |
| | *Cyclanorbis senegalensis* | 11300 | Gramentz 2008[d] |
| | *Cycloderma frenatum* | 14591 | Amniote Database |
| | *Lissemys punctata* | 1444.5 | Amniote Database |
| | *Lissemys scutata* | 1444.5 | Amniote Database |
| | *Nilssonia gangetica* | 19000 | Amniote Database |
| | *Palea steindachneri* | 10100 | Regis and Meik 2017 |
| | *Pelochelys bibroni* | 120000 | Bonin et al. 2006 |
| | *Pelodiscus sinensis* | 2327.5 | Amniote Database |
| | *Trionyx triunguis* | 10818 | Amniote Database |

a. Previously *G. puchra* was considered a subspecies and both reach similar sizes.
b. *C. ciclormata* was consided the same species.
c. According to López-Luna et al. 2018, the morphology is similar to *K. scorpiodes.*
d. Data of. *C. elegans,* which reach similar size

##### *Phylogenetic signal*

To explore the phylogenetic signal of each trait, we estimated Pagel's $\lambda$ for each trait separately. This approach describes the strength of phylogenetic relationships on trait evolution under a Brownian motion model (Freckleton 2000). Pagel's $\lambda$ ranges between 0, which indicates that the patterns in the traits cannot be explained by the employed phylogeny, and 1, which indicates that the observed patterns in traits are tightly correlated with the placement of species in the phylogeny. We used the function *phylosig* in the package *Phytools* (Revell, 2012) to estimate the phylogenetic signal of each trait before the imputation analyses (Table S3).

**Table S3. Most life history traits in Testudines and Crocodilia species have a strong phylogenetic signal.** Pagel's $\lambda$ describes the statistical dependence among species' trait values due to their phylogenetic relationships, ranging between 1 (meaning a life history trait is fully related to the phylogenetic structure as explained by Brownian motion), and 0 (meaning no phylogenetic structuring of the trait). Pagel's $\lambda$ was calculated for all traits included in the analyses. *N* represents the number of species of Testudines and Crocodilia with data for each trait.

| Life history trait | Symbol | *N* | Pagel's $\lambda$ | |
|---|---|---|---|---|
| Adult survival | *Sa* | 35 | ~0.001 | [0.00 – 0.32] |
| Juvenile survival | *Sj* | 33 | 0.310 | [0.00 – 0.81] |
| Age at sexual maturity | *Lα* | 114 | ~0.001 | [0.00 – 0.99] |
| Clutch size | *CS* | 253 | 0.990 | [0.97 – 0.99] |
| Mean number of clutches per year | *CN* | 98 | 0.920 | [0.77 – 0.97] |
| Maximum lifespan | *ML* | 207 | 0.820 | [0.65 – 0.91] |
